# Regression Based Accuracy Estimation for Multiple Sequence Alignment

**DOI:** 10.1101/2022.05.22.493004

**Authors:** Luis Cedillo, Hector Richart Ruiz, Dan DeBlasio

**Affiliations:** Computer Science Department, University of Texas at El Paso

## Abstract

Multiple sequence alignment plays an important role in many important analyses. However, aligning multiple biological sequences is a complex task, thus many tools have been developed to align sequences under a biologically-inspired objective function. But these tools require a user-defined parameter vector, which if chosen incorrectly, can greatly impact down-stream analysis. Parameter Advising addresses this challenge of selecting input-specific parameter vectors by comparing alignments produced by a carefully constructed set of parameter configurations. In an ideal scenario, we would rank alignments based on their accuracy. However, in practice, we do not have a reference from which to calculate accuracy. Therefore, it becomes necessary to *estimate* the accuracy to rank the alignments. One solution involves the use of estimators such as `Facet`. The accuracy estimator `Facet` computes an estimate of accuracy as a linear combination of efficiently-computable feature functions. In this work we introduce two new estimators called `Lead` (short for <u>L</u>earned <u>a</u>ccuracy <u>e</u>stimator from large <u>d</u>atasets) which use the same underlying feature functions as `Facet` but are built on top of highly efficient machine learning protocols, allowing us to take advantage of a larger training corpus.

**Note about previous versions:** A previous version of this paper was released on bioRxiv and presented the results of our previous study (`Facet`) with an error. This error has been corrected, and the conclusions made have been updated based on this new data. This corrected version stands as reference for anyone who may have encountered the versions with inaccuracies.

## 1 Introduction

For many computational biologists, aligning multiple biological sequences, mainly protein sequences, is an essential step in analysis. These alignments provide key insights into the correlation of various regions in the protein, and can be a guide to determining evolutionary history. From a computational standpoint though the problem is complicated, and in fact, it was proven that finding an optimal alignment of a set of sequences is NP-Complete [31, 16]. Because of this juxtaposition of the importance and complexity of the problem there are many existing heuristics that find *good* (though not optimal) multiple sequence alignments [27, 26, 13, 10, 22, 32, among others]

None of these tools address a fundamental issue: the multiple sequence alignment problem requires as part of its input a user-defined objective function. While there are several well-studied objectives, the most common of which is sum-of-pairs, this still leaves the open question of the function’s parameters: the rewards and penalties associated with each operation used to construct the alignment. Thus when new users, or even experienced users with a new class of inputs, approach the task of creating a multiple sequence alignment, they are required to choose a value for each of the tunable parameters of the tool (we call the collection of parameter values used is called a *parameter vector*). Making the *wrong* choice can have a highly detrimental impact on downstream analysis; manually tuning these algorithm values in a systematic manner requires not only a more-than-cursory understanding of the underlying application and the data, but also substantial amounts of time. Because of this most users rely upon the *default* parameter vector included with each tool. These defaults are specifically set to provide good results on average across all types of input, but the most interesting biological datasets are typically far from average.

Most alignment objective functions, and in turn the tools that utilize them, have parameters for both the penalty assessed for adding gaps into a sequence and the penalty/reward for the substitution (match/mismatch) of any two amino acids (residue). This means for the 20-character amino acid alphabet used for protein sequences, 190 individual values must be specified by the user for substitutions alone. Thankfully this is a well-studied problem, and there are many highly-accurate pre-calculated databases of substitution scores, called *substitution matrices*. Typically users are given a choice from a class of matrices like `VTML` [21], `BLOSUM` [12], or `PAM` [6]. Within each class, there are different matrices available that are optimized for sequences with a specific level of entropy. When substitution matrices are used, the total number of tunable parameters is greatly reduced, simplifying (though not eliminating) the parameter choice problem.

One confounding factor for making an informed parameter vector choice for multiple sequence alignment is that unlike some domains, such as transcript assembly which has a readily available quality measurement which can be used on any input [9], the standard method for assessing accuracy requires comparison to a reference (ground-truth) alignment. Thus, the accuracy of a computed alignment can only be measured on so-called *benchmark* sequence sets. Typically these benchmarks are constructed by aligning the three-dimensional structure of proteins to find amino acids that are highly correlated. Those groups of amino acids (one from each protein) that are within some threshold of distance in the 3D alignment are turned into complete columns of the sequence alignment. These anchoring positions are called *core columns*. Accuracy of a computed alignment is then calculated as the fraction of substitutions from the core columns of the reference alignment that are recovered. Several databases of these benchmarks exist and are regularly used to compare multiple sequence alignment tools [3, 11, 25, 29, 2].

When a new (non-benchmark) sequence set is presented and aligned, there is no reference alignment from which to measure accuracy. In this case, one is left to estimate the accuracy of the computed alignment. There are a variety of tools available to compute this estimation, which can generally be divided into two main categories: *scoring-function-based* tools [14, 4, 28, 1] which calculate a value based on the combination of measurable attributes of the alignment alone; and *support-based* tools [19, 18, 24] which use a collection of alignments over the same sequences to label each one in the set.

Using one of the tools above to estimate accuracy, a user can then begin to search for the optimal parameter vector for their input. While this can be done by hand or using an off-the-shelf iterative optimization mechanism [such as coordinate ascent; 33], this is typically still quite time consuming. To automate this process DeBlasio and Kececioglu [8] developed the *Parameter Advising* framework, which without additional wall-clock time if the appropriate resources are available can make input-specific parameter choices for multiple sequence alignment (details of an advisor are contained in Section 2 to make this work fully self-contained). In that work, they show the advisor using the `Opal` alignment tool and the `Facet` accuracy estimator proved to have the highest advising accuracy.

In this study, we present a new estimator `Lead` (short for Learned accuracy estimator from large datasets) in two variants: `Lead-MLP` (which uses a multi-layer perceptron) and `Lead-LR` (which uses linear regression). These two new scoring-function-based accuracy estimators reimagine the `Facet` estimator by using modern machine learning techniques for optimization, rather than combinatorial optimization, to exploit the much larger datasets we have produced.

Although `Lead-MLP` and `Lead-LR` rely on the same underlying set of efficiently-computable feature functions as `Facet`, they stand out as better regressors as they show a much stronger correlation with true accuracy due to being able to optimize for this task specifically. This becomes evident when we look at their substantially higher R-squared (*R*^2^) scores both in training and testing.

While `Lead-MLP` and `Lead-LR` excel as better overall regressors with lower Mean Squared Error (MSE) and higher R-squared (*R*^2^) scores compared to `Facet`, it’s worth noting that both `Facet` and the new estimators perform quite similarly when it comes to parameter advising.

## 2 Summary of Prior Work

Parameter Advising, depicted in Figure 1, takes the same input as the underlying scientific application and returns a single result. So to the end-user it appears to be just another tool to solve an existing problem but in this case a tool without any tunable parameters. But the process of selecting a parameter vector that provides higher-than-default accuracy, tailored to user input sequences, is abstracted away.

**Figure 1:**
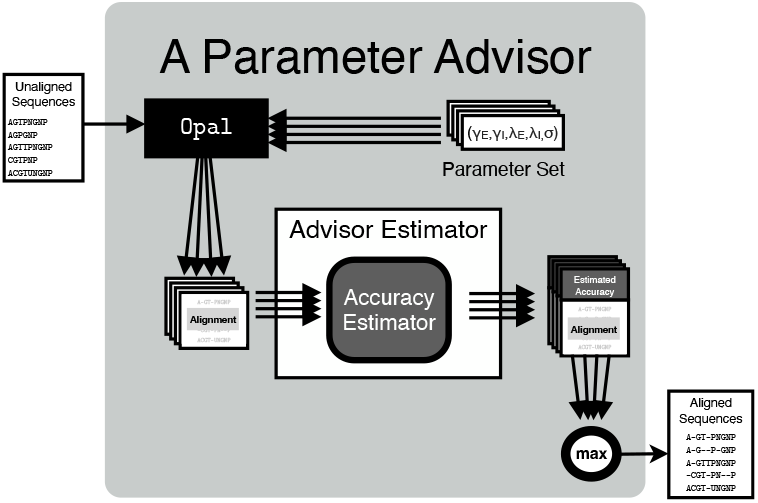
The parameter advising process.

An advisor contains two key components: a set of candidate parameter vectors, called an *advisor set* ; and an accuracy estimation tool used to choose from among those vectors, called an *advisor estimator*. The actual construction of these two components is described in detail below.

The Parameter Advising procedure is as follows:

- The input is combined with each of the parameter vectors in the advisor set (and the underlying application) to create a set of *candidate output* (alignments).
- The advisor estimator is then used to label each of the candidate alignments with an estimated accuracy.
- Finally, using the labeled estimated accuracy the candidate alignment that is predicted to be most accurate is returned to the user, therefore selecting the parameter vector.

Parameter advising is an *a posteriori* selection method and may not work in all cases, but in those applications where resources can be allocated such that each of the candidate outputs and their labels can be produced in parallel, the advising procedure adds only a negligible amount of wall-clock time (the time needed to run the estimator, also in parallel, and select the best output). In addition to multiple sequence alignment, this framework has been shown to work well for selecting parameters for transcript assembly [9].

Throughout the paper it is assumed that there is a consistent set *B* of benchmark sequence sets (the precise sets themselves will be explained in Section 3.3). Because there is bias in the sampling of these benchmarks (as they are made by hand, additional commentary on this in later sections), it is assumed that at all stages each benchmark will have a weight (or importance), *w*_*b*_ for *b ∈ B*. This weighting will be used to correct for the overabundance of some types of examples.

### 2.1 Advisor Sets

For the task of parameter advising an advisor set need to satisfy two main criteria: the set should be small to reduce resource consumption, and should contain at least one parameter vector that will work well on each given input. Due to the somewhat contradictory nature of these two ideas, a method is needed to determine the balance between competing interests. One way to do this is to put an actual limit on the resource consumption by specifying *k*, the number of parameter vectors to include in our set. Experiments can be used to determine the cardinality at which there are diminishing returns, meaning adding more computational power does not provide substantial gains in advising accuracy.

Existing methods have been developed to accomplish the task of finding such sets with restricted cardinality. They all rely on first enumerating the entire universe *U* of parameter vectors for a tool across the entire set of benchmarks *B*. The combination of a benchmark *b ∈ B* and a parameter from the universe *p ∈ U* produces an alignment, 𝔸_*bp*_. Because the reference alignment is known the true accuracy, *A*_*bp*_, can be calculated. If there is a pre-existing accuracy estimator it is assumed that the estimated accuracy for each of these alignments is *E*_*bp*_.

The *oracle* set finding method uses only the true accuracy, and finds sets that are optimal if you had an estimator function that predicted accuracy exactly (an oracle). Finding optimal sets while taking the accuracy estimator into account is not practical, thus a *greedy approximation* method was developed that works well in practice.

#### 2.1.1 Oracle Sets

While finding optimal parameter sets is NP-Complete [7] whether you have the estimator values or not. For a small enough universe the Oracle set problem can be solved exactly using an integer linear program (ILP).^1^ The program contains variables *s*_*p*_ which indicates if parameter *p* is in the advisor set, and variables *v*_*bp*_ that will signify if the advisor using the found set chooses parameter *p* for benchmark *b*. Three groups of constraints then ensure that:

1. for each benchmark has only one chosen alignment,

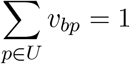
2. an alignment is only chosen if the associated parameter is chosen,

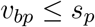

and (3) that only *k* parameters are chosen.

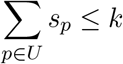

With the constraints in place, the objective then would be to maximize the total accuracy of all of the chosen alignments

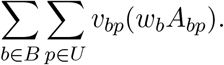

The ILP then contains *O*(|*U* ||*B*|) variables and constraints, though only *O*(|*U* |) of the variables need to be explicitly declared as binary (integer), if the parameter variables are binary the alignment variables must be binary valued for a solution to be optimal (with the exception of ties in accuracy, but this would not impact optimality).

#### 2.1.2 Greedy Sets

Finding optimal estimator-aware sets for large cardinalities is impractical, but a greedy procedure has been shown to provide substantial accuracy increases (over Oracle sets) in practice. Assume that an advisor *𝒜*_*δ*_(*P*) takes as input the set of parameter vectors *P ⊆ U*, and returns the average advising accuracy (over all benchmarks in *B*) of a parameter advisor that uses *P* as its advisor set (it has access to the set of benchmarks, true accuracies, and estimated accuracies). For generalization purposes during training (but not for showing the results in later sections), a margin of error on the estimator, *δ*, is used within the function *𝒜*. Rather than using a single alignment’s accuracy for a benchmark (the one with highest estimated accuracy), the method calculates the average for all alignments of a benchmark that are within *δ* of the maximum [details of this can be found in 7]. The initial set *P*_1_ is the Oracle set of size 1, in other words the best single default parameter vector. At each step the greedy procedure finds the *next best* parameter to add that is not already in the set, i.e. the one that when added provides the highest advisor accuracy. That is to say the greedy method at iteration *i* first finds

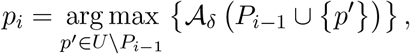

then sets *P*_*i*_ = *P*_*i−*1_ *∪ {p*_*i*_*}*. It continues with this procedure until *P*_*k*_ is found.

### 2.2 Advisor Estimator

It is the advisor estimator’s responsibility to rank the candidate outputs in such a way that the highest-ranked alignment is the most accurate. The best estimator is referred to as the Oracle (also used above) which knows the true accuracy and would be able to make a ranking based on this information. In the absence of an Oracle something close needs to be found. There are in theory two types of advisor “estimators”: one that assigns an estimated accuracy to each candidate (as will be used here), and another that simply chooses from the set of candidates. This second type of advisor is yet to be explored, but would still likely need to rely on some way to extract information (features) from the alignment itself in order to make a judgment.

#### 2.2.1 Feature Functions

The feature functions used were heavily studied in previous work, they consist of efficiently-computable functions that extract a singular numeric value that is correlated with accuracy. This set contains both canonical feature functions that have been historically used for benchmarking as well as some developed specifically for this task. Because the focus is on *protein* multiple sequence alignments, many of the feature functions will exploit the predicted secondary structure of the sequences in the alignment to calculate a value. This means each amino acid in the alignment is labeled with both a class (*α*-helix, *β*-strand, or coil) and a set of three probabilities which the features utilize.

The features developed are also non-local, meaning they generally use information from across the entire alignment. This means that they contain more information than an alignment objective function (which by design are local calculations).

Details of the feature computations can be found in Kececioglu and DeBlasio [14], but they are listed here for convenience as they are a crucial component of our new methods, roughly in order of importance:

- **Secondary Structure Blockiness** — maximum percentage of an alignment that can be covered by a packing of contiguous two-dimensional blocks of residues with the same secondary structure class label.
- **Percent Identity** and **Secondary Structure Percent Identity** — the fraction of aligned amino acids (structure class labels) in all extracted pairwise alignments from the input that are precisely the same.
- **Average Replacement Score** — average scaled BLOSUM62 score for all aligned amino acids from the induced pairwise alignments.
- **Gap Extension Percentage** and **Gap Open Percentage** — percentage of characters in the alignment that are (runs of) gap characters.
- **Information Content** — average over all columns of the difference in distribution of amino acid frequencies compared to a background distribution.
- **Substitution Compatibility** and **Gap Compatibility** — fraction of pairs of columns that pass the 4 gametes test when converted into 1 and 0 by use of majority/minority amino acid character or gap/non-gap character.
- **Structure Agreement** — weighted sum of structure class probabilities for a window around each aligned pair of amino acids in induced pairwise alignments.
- **Gap/Coil Density** — fraction of gap characters that align with the coil structure class label.
- **Core Column Density** — fraction of columns that are gap-free and contain mostly the same amino acid.
- **Sequence Consensus** and **Gap Consensus** — fraction of pairs of columns that have the same majority/minority amino acid character or gap/non-gap character pattern.

All feature values are normalized to be in the range [0, 1], and if the value cannot be defined (i.e. gap open percentage when there are no gaps) the value is fixed at 0.

#### 2.2.2 The Facet Estimator

The `Facet` (short for <u>F</u>eature-based Accuracy <u>E</u>s<u>t</u>imator) model uses a linear combination of the feature values to calculate an accuracy estimate. Given a set of weights *T* = (*t*_1_, *t*_2_, …, *t*_*n*_) the `Facet` value is computed from a set of feature functions *F*_1_, *F*_2_, …, *F*_*n*_ as

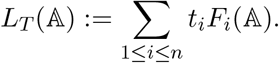

There are several methods that can be used to learn an accuracy estimator, one can either try to match the estimator values to the true accuracy exactly or, what worked better in previous experiments for the task of parameter advising for multiple sequence alignment, focus on the objective of ranking candidates.

All of the `Facet` results shown will use the methods described in DeBlasio and Kececioglu [8], which learns the values of *T* using combinatorial optimization (in this case linear programming, LP). In this case for all pairs of alignments 𝔸_*bp*_ and 𝔸_*bq*_ aligning the same benchmark using parameters *p* and *q* such that *A*_*bp*_ *> A*_*bq*_, the goal is to minimize the ranking error:

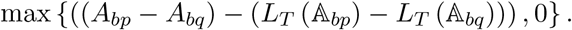

Note that in this case, the only free variables are *T*, because the alignments are fixed then the feature values and true accuracies are also. Additional constraints are added to ensure that the estimator as well as the values in *T* are in the range [0, 1].

## 3 Methods

### 3.1 Lead-MLP: Exploiting Non-Linearity

#### 3.1.1 Learning hyperparameters

To address the regression challenge we discussed earlier, we employed two different models: a multilayer perceptron (MLP) and a linear regressor (LR). These models were trained using efficient machine learning techniques and a significantly larger training dataset.

While some neural network architectures can be inspired by the underlying structure of the input data [20], in situations where this is not feasible, we rely on guided search to find the most appropriate architecture. Training a network with an excessive number of parameters can be time-consuming and resource-intensive. Furthermore, given the substantial size of our dataset, deeper neural networks have a tendency to overfit, which can hurt their ability to generalize effectively. The key is to strike the right balance between accuracy, complexity, computation resources, and generalization As a result, we excluded certain architecture configurations from our search space, as is common practice in most studies. Our experiments utilized a nested cross-validation approach to identify the optimal neural network architecture, as depicted in Figure 2. In a cross-validation approach training samples are initially split into *k* partitions, and in each of the *k* top-level instances one of the partitions is held out for testing, and the other are in the training set. In the nested case, a lower level split is made of the remaining *k −* 1 partitions with one of these being used for *validation* and the other *k −* 2 being used for testing.

**Figure 2:**
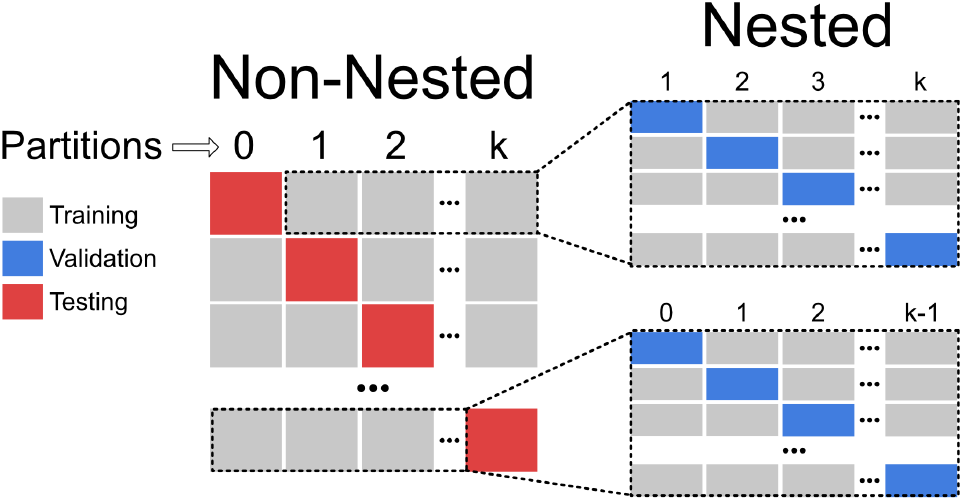
Visual depictions of both cross-validation mechanisms.

We generated a set of 64 unique architectures through randomization, aiming for a balance between performance and computational demands. These architectures were intentionally kept compact, comprising of a maximum of two hidden layers and with a maximum of 16 neurons in each layer.

Our local search algorithm explored these architectures iteratively and selected the one that minimized the average mean squared error on all *k –* 1 validation sets. During the search, we implemented early stopping criteria during training. This enabled us to stop training architectures that were not showing promising results on the validation set.

Figure 3 shows the architecture that performed the best on 9 out of the 14 folds.

**Figure 3:**
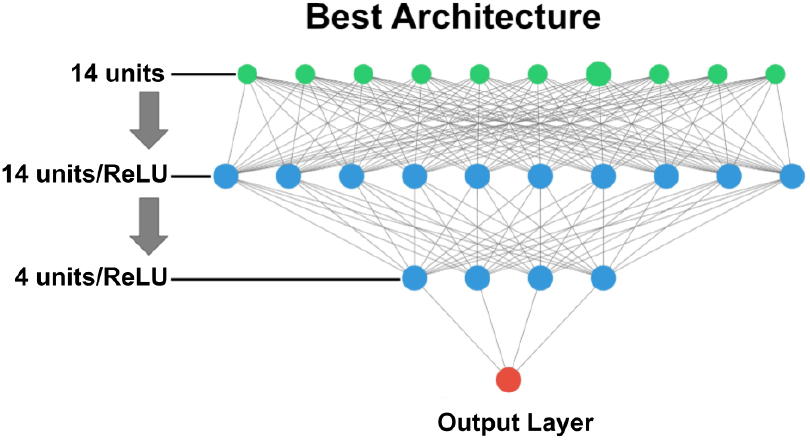
Visual depiction of the model architecture that was optimal for a majority of cross-validation folds.

#### 3.1.2 Training the model

After identifying an architecture it was time to train the final model and compare its final performance with unseen data. For this step, we took the architecture that displayed the best performance in each fold and trained the final model on the entire training set of that respective cross-validation fold.

`Lead-MLP` was implemented using the `MLPRegressor` of `scikits-learn`.

All models were trained with a batch size set to “auto”, employing the default learning rate of 0.001. The ReLU function served as the activation function, and we used the Adam optimizer. Additionally, the Mean Squared Error was chosen as the loss function.

It’s worth noting that the *α* (set to 0.1) hyper-parameter played a crucial role in controlling the strength of the *L*_2_ regularization term applied during training. When regularization is applied, a penalty term is added to the cost function to prevent overfitting.

All results are reported as an average over all of the folds.

#### A note about implementation

All of the methods above as well as the experiments below were performed using `scikits-learn` [23]. All models, architectures, and training scripts (written for `python3`) for both `Lead-MLP` and `Lead-LR` are released on the DeBlasio Lab GitHub, at `github. com/deblasiolab/Lead`.

### 3.2 Lead-LR: Utilizing More Data

As mentioned in Section 2.2.2, `Facet` was trained to minimize the ranking error of the accuracy prediction made by the tool. This was done mainly because it showed better performance than trying to match the accuracy value directly. One reason this was likely the case is that the LP method used was not able to scale to large numbers of samples practically, and thus narrowing the task to the actual use was better with limited data. But, modern machine learning optimization tools now allow us to use much more data to train models, they also enable us to fully utilize complex hardware to accomplish this task.

`Lead-LR` was implemented using the Linear Regression model of `scikits-learn`. A separate estimator was learned on the training set of each cross-validation fold, minimizing the residual sum of squares between the predicted and true accuracy of each alignment. All results are reported as an average over all of the folds.

The weights given by `Lead-LR` trained on a single fold are in Table 1, sorted by the absolute value of their influence. Note that some of the features have negative weights, this was not allowed in `Facet`. While not intuitive, since all of the features originally trended with accuracy, it can be explained: for instance if an alignment has a high sequence identity but also a high gap density, these two values (because they have opposite signs) will counteract each other. More of the features in `Lead-LR` have significant weight compared with `Facet`, in the previous studies only 5 features ended up being used while here more than 9 features make contributions to the final score.

**Table 1:**
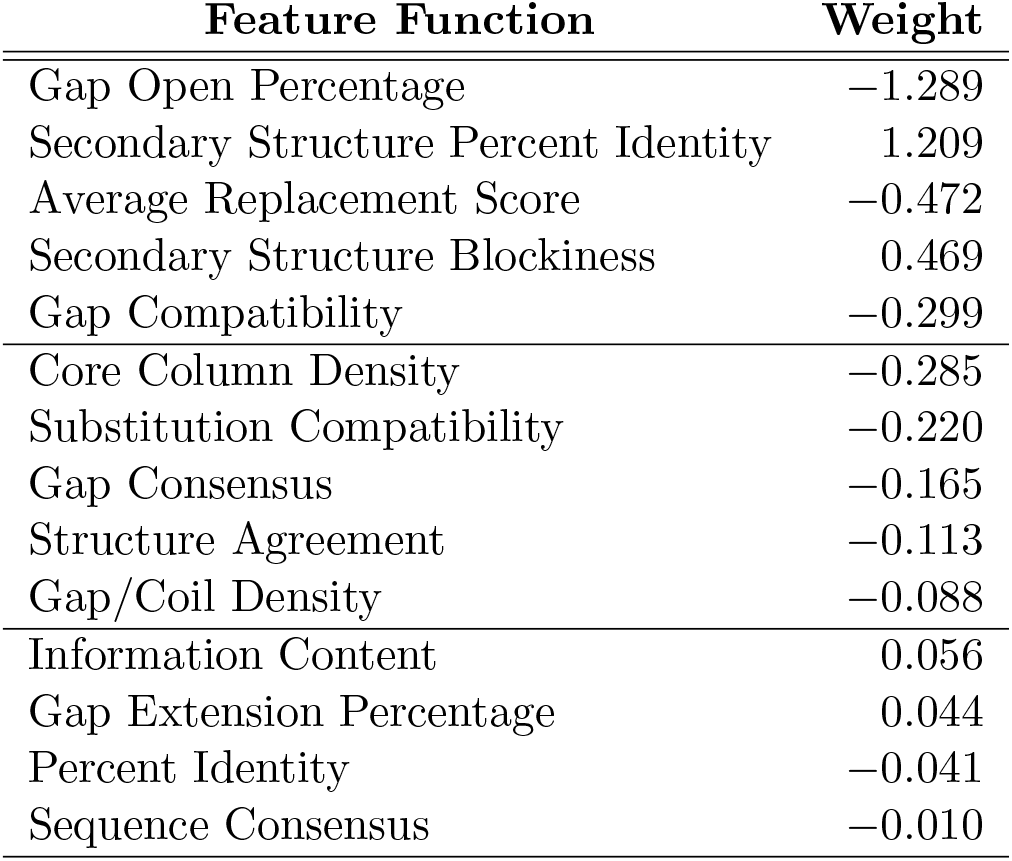
Example Lead-LR Feature Weights.

| Feature Function | Weight |
| --- | --- |
| Gap Open Percentage | -1.289 |
| Secondary Structure Percent Identity | 1.209 |
| Average Replacement Score | -0.472 |
| Secondary Structure Blockiness | 0.469 |
| Gap Compatibility | -0.299 |
| Core Column Density | -0.285 |
| Substitution Compatibility | -0.220 |
| Gap Consensus | -0.165 |
| Structure Agreement | -0.113 |
| Gap/Coil Density | -0.088 |
| Information Content | 0.056 |
| Gap Extension Percentage | 0.044 |
| Percent Identity | -0.041 |
| Sequence Consensus | -0.010 |

### 3.3 An Expanded Parameter Universe

Two *universes* of parameter vectors are used: one containing just over 2000 parameter vectors used for `Facet` called “original”, and an extended set of 16,896 parameter vectors called the “extended” universe. The original set was used to construct all of the Oracle advisor sets as well as the `Facet` estimator, the extended set was used for training `Lead-MLP`, `Lead-LR`, and for the Greedy advisor sets.

The extended set was originally developed for use in the `Facet` methodology. It is constructed by first selecting 8 commonly use high-accuracy replacement matrices (`VTML20`, `40`, `80`, `120`, `200`, and `BLOSUM45`, `62`, `80`). For each of the four `Opal` gap penalty parameters (gap-open and gap-extension for both terminal and non-terminal gaps) several possible values were enumerated, and the cross product of these discrete values and the replacement matrices produced a collection (universe) of parameter vectors. Using the default parameter vector as a starting [which was found via inverse parametric alignment; 17, 15] we sampled up to five additional values for each parameter above and below the default. In the case of the terminal gaps, a value relative to the corresponding non-terminal parameter value was chosen rather than a specific penalty. This set constituted approximately 2,000 choices of gap penalty combinations, leading to the number of parameter vectors above in our extended universe.

The original universe is then an informed subsampling of this universe, used to allow the older methods to be solved.

Building on the datasets used in DeBlasio and Kececioglu [8], a carefully curated set of 861 benchmark sequence sets is used consisting of examples from PALI [3] and BENCH [11]; which is itself a collection of alignments from OxBench [25], SABRE [29], and BAliBase [2]. Each benchmark consists of a set of protein sequences as well as a reference alignment which is generally induced using an alignment of the known underlying three-dimensional protein structures. The alignment information is removed to also retrieve the set of constituent sequences. Because the reference is known, the accuracy can be determined as defined earlier.

As mentioned in Section 2 the set of benchmarks is biased toward easy-to-align sets of sequences. This is likely because these benchmark alignments are constructed mostly by hand, and thus only the alignments with which they have the most confidence end up being released. But, in practice, it is expected that sequence sets from across all levels of difficulty will be input. To correct this, each benchmark’s ease of alignment was classified using its accuracy using the `Opal` aligner’s default parameter vector. This parameter vector was chosen specifically to work best *on average*, and it is the set of parameter values most users will use. The range of accuracies was then divided into 10 equally sized bins and assigned a benchmark to the bin that corresponds to its accuracy when aligned using the default parameter vector. To exemplify the bias in the sample, note that there is a more than 30x increase in the size of the easiest-to-align bin (those with accuracy above 90%, 431) and the hardest-to-align (those *≤* 10%, 14). Unless otherwise noted, the weights *w*_*b*_ will be assigned to evenly distribute among *bins* rather than benchmarks. This means with this scheme the accuracy of using only the default parameter vector will be close to 50%, rather than near 80% when weighting by benchmark.

Due to the binning described above, this number of cross validation partitions is somewhat forced since the smallest bins have 14 benchmarks a piece. In each case, each bin was divided randomly into 14 partitions as described earlier. All of the results below show the average across all 14 cross-validation folds.

## 4 Experimental Results

### 4.1 Accuracy of the Estimator

Figure 4 shows the comparison of the estimated value (vertical) with the true accuracy (horizontal) for each of the 14 million alignments in the dataset across the two new estimators and `Facet`. Each alignment is shown for when the benchmark it is aligning is in the testing set. To work well as an advisor estimator the value should trend with true accuracy and the plot should have high slope and low spread in order to correctly distinguish between candidates. `Lead-MLP` and `Lead-LR` have both a steeper slope and less spread than `Facet`. The trend-line is shown that is the best linear fit to all of the alignments plotted. The conclusion that `Lead-MLP` and `Lead-LR` outperform `Facet` on the task of matching accuracy is also corroborated by the R-squared values shown, `Lead-MLP` has a value of 0.744 and `Lead-LR` 0.684 while `Facet` is only 0.413.

**Figure 4:**
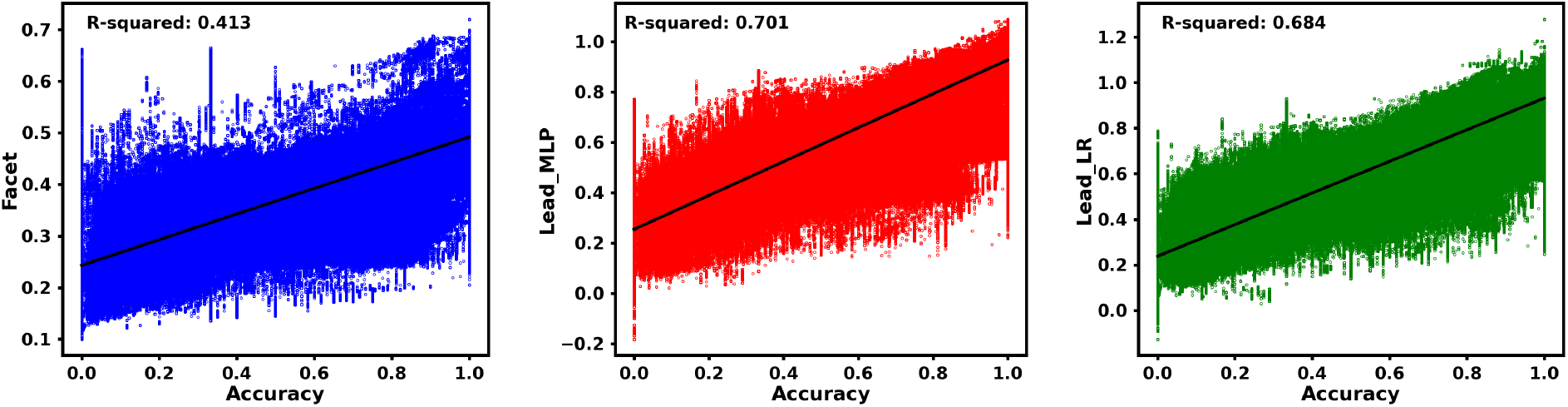
Estimated versus true accuracy of all alignments in the dataset for `Facet`, `Lead-MLP`, and `Lead-LR`.

One contributing factor to the difference seen in the correlation of the estimator and accuracy is that due to the computational limitations at the time, the `Facet` estimator is learned over only parameter pairs in the Oracle set of a given size (in the case of the results above cardinality 25), while because `Lead-MLP` and `Lead-LR` are able to take advantage of modern GPUs, the whole training set was able to be exploited. This change in the training set used in `Facet` also means there is variation in the learned weights at different cardinalities as well as between folds (which will be shown in later sections). As mentioned in Section 3.1 unlike `Facet`, `Lead-MLP` and `Lead-LR` were trained to match the accuracy of a given alignment exactly. Therefore, the objective of `Lead-MLP` and `Lead-LR` estimators is toward increasing R-Squared.

One other item to note in Figure 4 is the scale on the 3 plots. Because there is nothing tying the estimator values of `Facet` to the true accuracies the range of values ends up being quite small. In contrast, by training to the actual accuracy we see estimated accuracies across the whole range from 0 to 1; but this comes at the expense of having predicted values that are somewhat contrary to the definition of accuracy, as you cannot recover fewer than 0 (or more than all) pairs of residues.

### 4.2 Accuracy of the Advisor

The main purpose of developing `Lead-MLP` and `Lead-LR` was to improve accuracy of the resulting parameter advisor. In the experiments below `Lead-MLP`, `Lead-LR`, and `Facet` are used as the accuracy estimator to construct several advisors. Advisor-oblivious Oracle sets (using the original universe) and advisor-aware Greedy sets (using the extended universe) are used in combination with the two advisors to create 4 classes of advisors. The `Facet` method relies on a parameter vector set to learn its weights, thus, in all experiments, the `Facet` estimator used was trained on the Oracle set of the specified cardinality, whereas only a single `Lead-MLP` and `Lead-LR` advisor is used for each cross-validation fold.

#### 4.2.1 Oracle Sets

Figure 5 shows the accuracy of the three advisors using Oracle sets on the original universe of parameter vectors as well as accuracy of the default parameter vector. The two plots show the use of the advisor averaged across the 14 cross-validation folds’ training (right) and testing (left) benchmarks. Each point on the four lines represents one advisor with its position corresponding to the advisor set cardinality (horizontal) and the advising accuracy (vertical).

**Figure 5:**
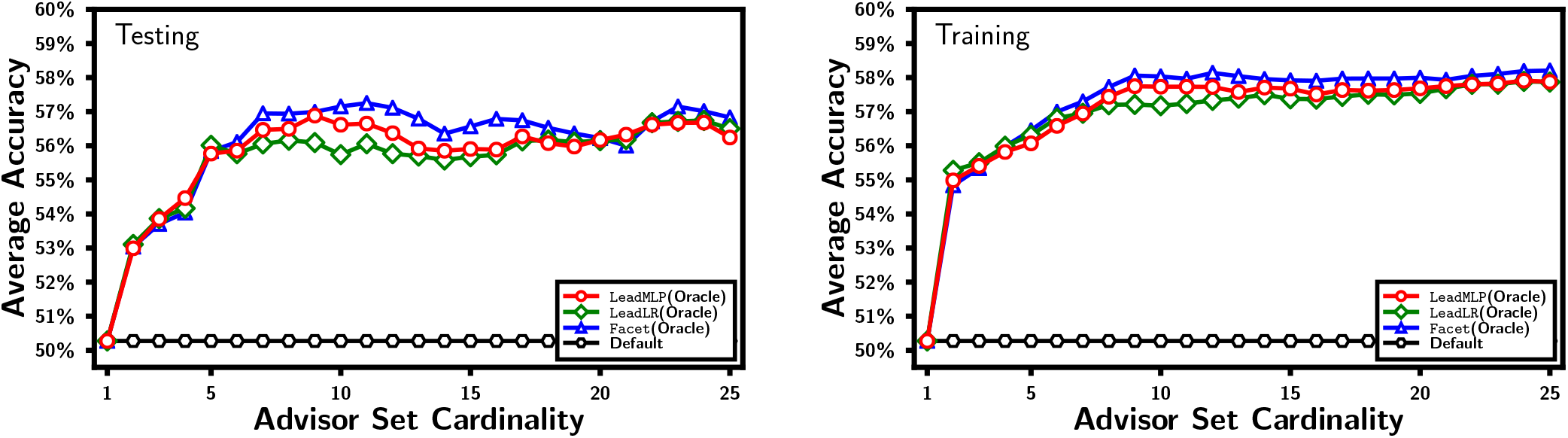
Comparison of the advising accuracy of `Facet`, `Lead-MLP`, and `Lead-LR` using Oracle advisor sets

#### 4.2.2 Greedy Sets

Figure 6 shows the accuracy of advisors using Greedy advisor sets trained specifically for the individual estimators. Once again, the two plots show the use of the advisor averaged across the 14 cross-validation folds’ training (right) and testing (left) benchmarks. Just as above, each point on the four lines represents one advisor with its position corresponding to the advisor set cardinality (horizontal) and the advising accuracy (vertical). While we tested various values of *δ* [as was done in 8], we found *δ* = 0 performed best and this was used to produce all figures.

**Figure 6:**
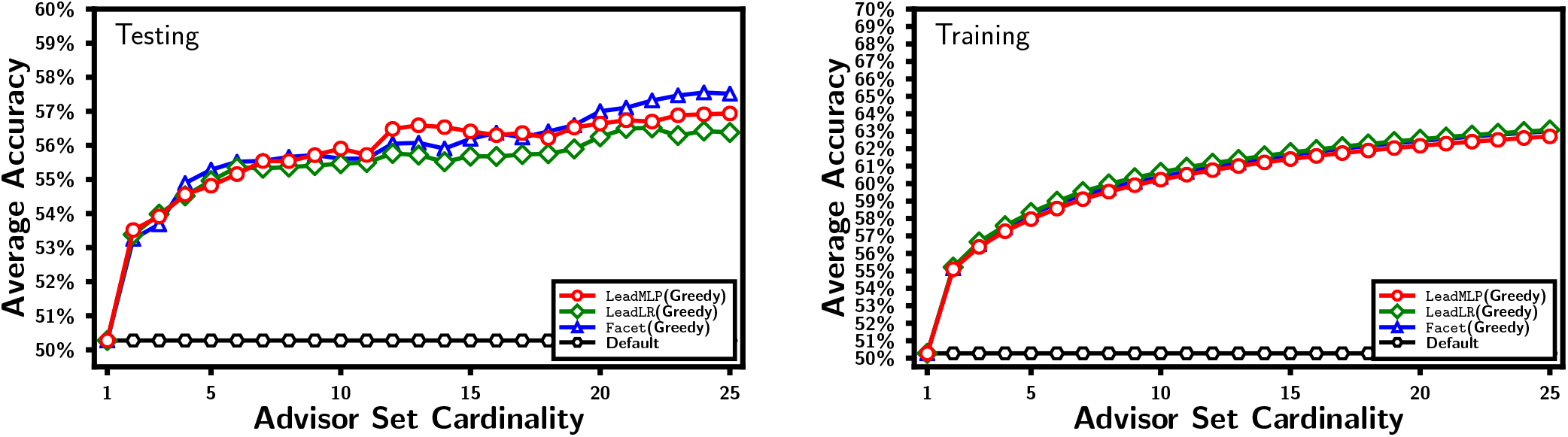
Comparison of the advising accuracy of `Facet` and `Lead-MLP` using Greedy advisor sets

`Lead-MLP` shows consistently higher advising accuracy than the two linear estimators on all but the smallest advisor sets. `Lead-MLP` nor `Lead-LR` benefit as much from Greedy sets as `Facet` with respect to the training benchmarks, seeing an average increase of 3.0% and 3.4% compared to using Oracle sets respectively. While at small cardinalities `Lead-MLP`, `Lead-LR`, and `Facet` show similar performance, across the cardinalities shown the average increase in advising accuracy is still 2.2%. This is mainly due to the fact that the `Facet` estimator is inconsistent in its accuracy as you vary the set cardinality; while `Lead-MLP` and `Lead-LR` continue to improve as the carnality rises.

### 4.3 Comparison of Alternate Machine Learning Models

Choosing the right algorithm to train a model is crucial. To evaluate the performance of our neural network, we trained models using both decision trees and random forest regressors optimized to minimize squared error. To further improve their performance, we also conducted a local search to find the optimal value for the maximum depth hyperparameter for both algorithms. The maximum depth values tested were 5, 10, 25, 35, and 50. Both the random forest regressor and decision tree regressor converged to the max depth value of 35. The estimated accuracies of these models are shown in Supplemental Figure 1. Both `Lead-MLP` and `Lead-LR` have a higher correlation (R-squared) with true accuracy than any of the other models on the testing set. However, both random forest and decision trees showed an almost perfect correlation with the training data.

The estimators produced with the other machine learning methods were also tested as advisor estimators. The results are shown in Supplemental Figure 2. When this experiment is performed we see just how much the decision tree and random forest regressors overfit to the training benchmarks. On both the Oracle and Greedy advising sets, they have lower performance on the testing set than `Facet` (and in turn `Lead-MLP` and `Lead-LR`) while having training performances that are significantly higher than the other methods.

## 5 Discussion

In this study, we presented `Lead-MLP` and `Lead-LR`, two new scoring-function-based accuracy estimators that utilize the power of modern machine learning techniques for optimization, rather than combinatorial optimization, to exploit larger datasets.

One major issue with multiple sequence alignment tools is the need for user-defined parameter vectors, which can greatly impact downstream analysis if chosen incorrectly. Parameter Advising addresses this challenge by allowing for the selection of input-specific parameter vectors with high accuracy. However, to rank the available alignments, we need to know their true accuracy, but in practice, a reference from which to calculate this is not available. This is where our accuracy estimators, `Lead-MLP` and `Lead-LR`, come in. When used in Parameter Advising, our estimators show an increase of 6% over using only the default parameter vector.

We further implemented other machine learning methods to ensure that the choices of models presented were valid. The results seen using random forest and decision tree regressors can be found in the Supplemental Material.

For this specific regression problem, models with a large number of parameters, such as the random forest and decision tree, were prone to overfitting. These models, due to their complexity and excessive number of parameters, tended to fail to generalize to new, unseen data. On the other hand, models with fewer parameters, such as `Lead-MLP` and `Lead-LR` presented in this study, were able to perform well on both the training and unseen data given the limitations of the underlying benchmark sets available.

To keep this work succinct, we did not include results from other estimators such as TCS [4] or MOS [19] on the larger parameter universe of examples described here.

### 5.1 Conclusion

When estimating the accuracy of protein multiple sequence alignments a major improvement can be obtained by exploiting modern machine learning tools. It has been shown in previous studies that when estimating alignment accuracy, it is essential to extract features that help to create estimates for alignments of any size. For parameter advising, carefully constructed small sets of parameter vectors can be very effective, but using large amounts of data and models that can be trained on it shows even better accuracy using the same feature functions.

### 5.2 Ongoing Work

We avoided deeper MLP networks after finding that they were overfitting. This can occur when the model has too many parameters, making it able to memorize the training data but not generalize well to new data. On the other hand, shallow MLP networks have fewer parameters and are therefore less likely to overfit. In this specific case, the shallow MLP network was able to generalize better to unseen data and thus performed better on the regression problem. However we are currently exploring more advanced structures such as residual networks [5], and new multiple sequence alignment feature vectors that can be further exploited by deep learning architectures such as transformers [30].

While it should technically be possible, as formulated, finding Oracle Sets on the extended universe is currently intractable. As described in previous studies, the method uses an integer linear program, which was solvable in the original universe despite being technically NP-Complete. When constructed for the extended universe we were unable to find a solution within a week using either CPLEX or Gurobi (the two commonly used tools) on a machine with 256 threads and 2 terabytes of memory. Finding these sets would allow us to use the same universe at all stages of this study.

## Supporting information

Supplemental Graphs

## Acknowledgements

The authors thank Dr. Olac Fuentes and Jose Perez for fruitful discussions and the UTEP Campus Office of Undergraduate Research Initiatives (COURI). This work was partially funded by US Department of Education project number 226150864A, and the Computing Alliance of Hispanic-Serving Institutions (CAHSI) Research Experience for Undergraduates (REU) under NSF Grants CNS-2137791 and HRD-1834620.

## Footnotes

1 As is noted in Section 4 for the much larger universe used in most of the study is it not possible to solve these problems exactly at the time of publication so results are shown on the smaller parameter universe from previous studies for only the Oracle sets.

## Notes

https://github.com/deblasiolab/Lead

### Competing Interest Statement

The authors have declared no competing interest.

### Summary of Updates

Note about previous versions: A previous version of this paper was released on bioRxiv and presented the results of our previous study (Facet) with an error. This error has been corrected, and the conclusions made have been updated based on this new data. This corrected version stands as reference for anyone who may have encountered the versions with inaccuracies.

