## Supplemental Graphs for "Regression Based Accuracy Estimation for Multiple Sequence Alignment"

### Supplemental Figures

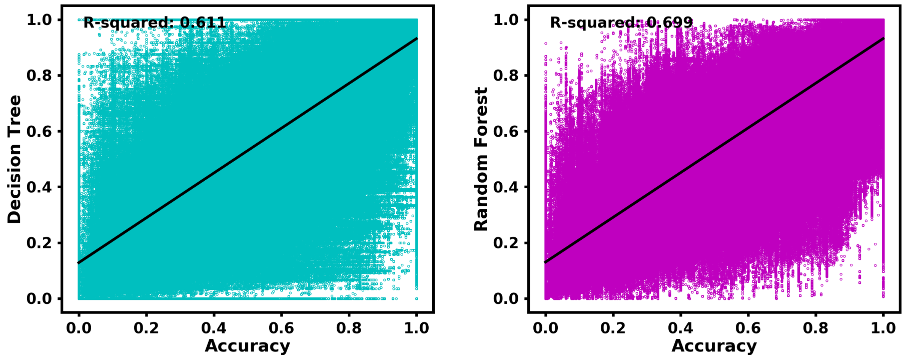


Supplemental Figure 1: Estimated versus true accuracy of all alignments in the dataset for other machine learning models.

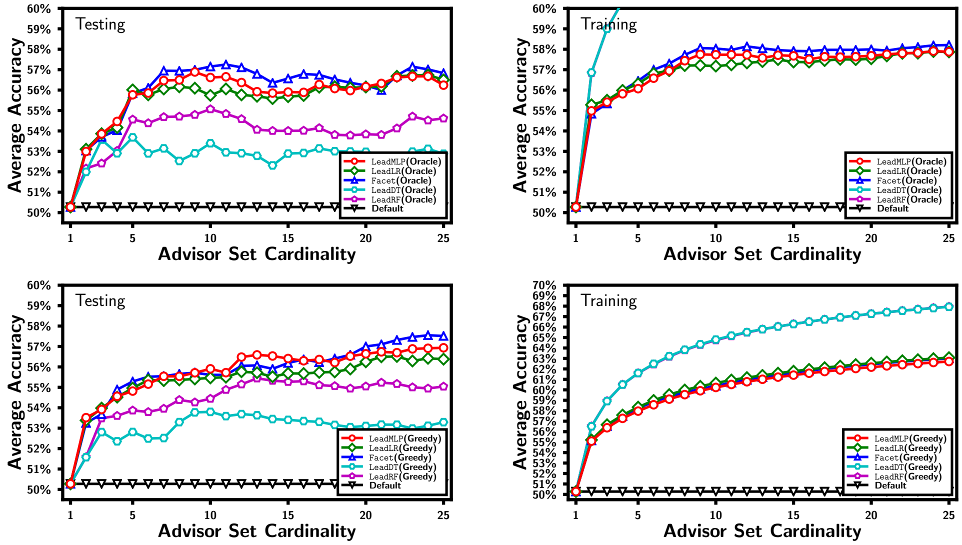


Supplemental Figure 2: Comparison of the advising accuracy of the other regressors
